# Early changes in the urine proteome in a rat liver tumor model

**DOI:** 10.1101/568246

**Authors:** Yameng Zhang, Yufei Gao, Youhe Gao

**Affiliations:** Department of Biochemistry and Molecular Biology, College of Life Sciences, Beijing Normal University, Gene Engineering Drug and Biotechnology Beijing Key Laboratory, Beijing 100875, China; College of Information Science and Technology, Beijing Normal University, Beijing 100875, China

**Keywords:** Urine, Proteomics, Liver tumor, Biomarkers

## Abstract

Urine, as a potential biomarker source among the body fluids, can accumulate many changes in the body due to the lack of a mechanism to maintain a homeostatic state. Previous studies have demonstrated that proteomic technology can find many potential biomarkers to reflect different diseases in the urine. This study aims to detect early changes in the urinary proteome in a rat liver tumor model. The tumor model was established with the Walker-256 carcinosarcoma cell line (W256). Compared to before the injection, ninety-five differential proteins were significantly changed in the experimental rats. At day 3, twelve proteins were identified in the absence of pathological changes, and four of them were altered at all four time-points (B2MG, VCAM1, HA11, and LG3BP). Seven had previously been associated with liver cancer. At day 5, fifty-two differential proteins were identified. At day 7 and day 11, there was a significant decrease in the body weight of the rats, and tumor tissue was observed in the liver. Fifty-two and forty differential proteins were changed significantly at day 7 and day 11, respectively. Of the proteins that were identified at these three time-points, and twenty-four were reported to be associated with liver cancer. Comparing the differential urinary proteins and biological processes of liver tumor model with those in different models of W256 grown in other organs, specific differential protein patterns were found among the four models, which indicates that the differential urinary proteins can reflect the differences when the same tumor cell grown in different organs.

**Significance:** This study demonstrated that (1) the rat liver tumor model caused early changes in urinary proteins may give new insight into the early diagnosis of liver cancer; (2) the same tumor cell grown in different organs can be reflected in differential urinary proteins.

## 1. Introduction

Liver cancer is the third leading cause of cancer mortality in the world, and the latest dataset on urban cancer in China also noted that liver cancer has one of the top three highest cancer mortalities^[1]^. It has the characteristics of substantial morbidity and mortality. The early detection of in situ or invasive carcinoma may prevent cancerous metastatic processes; therefore, it can significantly improve survival rates for cancer patients. Despite the technology of diagnosis to detect the cancer having advanced so quickly in the last decade, there are still many patients who cannot be diagnosed at early disease stages. To reduce mortality from cancer, novel approaches must be considered for early detection. One effective strategy to improve the prognosis of liver cancer is to find the tumor in the early stage, when the patients have no obvious symptoms so that liver function can be preserved as much as possible and more effective treatments can be applied. Early diagnosis and treatment are the most effective ways to prevent and treat cancer and reduce mortality^[2]^.

Currently, liver cancer diagnosis mainly relies on detection with imaging equipment (such as ultrasound, CT and MRI) and biomarkers; however, images are susceptible to operator experience, and it is difficult to distinguish between liver cancer and nonmalignant hyperplasia. It can also be hard to detect many small nodules in the early stage. Approximately 22% of early liver cancer imaging is not typical^[3]^. On the other hand, tumor biomarkers are used simpler to employ, but there are many challenges. For instance, alpha-fetoprotein (AFP), which rapidly decreases in serum after birth and is maintained at a low level throughout adulthood, is the most widely used biomarker for liver cancer^[4]^; however, serum AFP is not sufficient for diagnosis due to its poor sensitivity and specificity. Previous research suggests that no single serum biomarker can predict liver cancer with optimal sensitivity and specificity, particularly in the early stage^[5]^.

Urine is known to play an important role in body fluids and can reflect many changes in the body due to the lack of a mechanism to maintain a homeostatic state^[6]^. This is the main distinction between blood and urine. Many studies have demonstrated that proteomic technology can be used to find many potential biomarkers to reflect different diseases in the urine, such as glomerular diseases^[7]^, obstructive nephropathy^[8]^, hepatic fibrosis^[9]^, autoimmune myocarditis^[10]^, subcutaneous tumors^[11]^ and glioma^[12]^.

Animal models can help to us to better understand liver cancer and also have less complicated genetics, living and other conditions to monitor the development of the disease. In clinical studies, it is difficult to collect samples from a large number of liver cancer patients in the early stage. Before the onset of obvious symptoms, animal models can employ conditional control of the disease development process and provide more clues to find early biomarkers. This study used liquid chromatography coupled with tandem mass spectrometry (LC-MS/MS) to detect the urinary proteome in a rat liver tumor model. Several differential proteins associated with liver cancer were observed, and new evidence for biomarkers in the early stage was obtained.

## 2. Materials and methods

### 2.1 Animals

Male Wistar rats (130 ± 20 g) were purchased from Beijing Vital River Laboratory Animal Technology Co., Ltd. The animal license was SCXK (Beijing) 2016-0006. All experiments were approved by Institutional Animal Care Use & Welfare Committee of the Institute of Basic Medical Sciences, Peking Union Medical College (Animal Welfare Assurance Number: ACUC-A02-2014-007). All rats were housed under a standard 12-h light/12-h dark cycle, and the room temperature and humidity were maintained at a standard level (22 ± 1 °C, 65-70%).

### 2.2 Tumor cell line and Culture

Walker 256 tumor cells were cultured in the ascitic fluid of Wistar rats. All cells were harvested from the rats who were given an intraperitoneal injection of 107 Walker 256 carcinoma cells after two cycles of 7 d cell passage. Then, W256 cells were resuspended in phosphate-buffered saline (PBS) before injection. The viability of the cells was detected by the Trypan blue exclusion test using a Neubauer chamber.

### 2.3 Rat models of liver cancer

A tumor-bearing animal model was established in this study. All Wistar rats were randomly divided into different groups: the control group (n =5), and the Walker-256 (W256) tumor-bearing group (n =12). After anesthesia, the left medial lobe of the liver was exposed. W256 cells (2.5× 10^5^) were visually injected under the hepatic capsule into this lobe. The injection volume is 0.1 ml. An equal volume of PBS was also injected into the same location of the control rats.

### 2.4 Histological analysis

In the W256 model, the livers of the experimental and control rats were harvested 0, 3, 5, 7, 11 and 18 d after injection. For histopathology, the liver was fixed in formalin (4%) and embedded in paraffin. Then, all samples were sectioned and evaluated with hematoxylin and eosin (H&E) staining.

### 2.5 Urine collection and sample preparation

Urine samples of the W256 model were collected at four time-points, day 3, day 5, day 7 and day 11. Without any treatment, urine was collected from each rat by metabolic cage alone overnight. All rats were fasted and banned while collecting the urine sample. The urine samples were stored at -80°C for later use. Before LC-MS/MS analysis, the urine samples were thawed and transferred to centrifuge tubes for centrifugation at 12,000×g for 30 min at 4 °C to remove impurities. The samples were mixed with three volumes of prechilled ethanol, and the supernatants were precipitated at -20 °C for 2 h. The mixtures were centrifuged for 30 min at 4 °C, the supernatant was removed, and the precipitate was dissolved in a configured lysis buffer (8 mol/L urea, 2 mol/L thiourea, 50 mmol/L Tris, and 25 mmol/L DTT). After the dissolution was completed, the centrifugation was continued at 12,000×g for 30 min at 4 °C, then the supernatant was preserved. The protein concentration was determined by the Bradford assay. The urinary proteins at different time-points were digested using the FASP method^[13]^. One hundred micrograms of protein were added to the 10 kDa filter device (Pall, Port Washington, NY, USA) for each sample, and the protein was washed several times in sequence with the prepared UA (8 mol/L urea, 0.1 mol/L Tris-HCl, pH 8.5) and 25 mmol/L NH_4_HCO_3_ solutions. The protein samples were reduced with 20 mmol/L dithiothreitol (DTT, Sigma) at 37 °C for 1 h and then added to 50 mmol/L iodoacetamide (IAA, Sigma) for 30 min in the dark. The sample was centrifuged at 14,000×g for 30 min at 18 °C, then washed by UA and NH_4_HCO_3_, and trypsin (enzyme-to-protein ratio of 1:50) was added for overnight digestion at 37 °C. Oasis HLB cartridges (Waters, Milford, MA) were used to desalt the peptide mixtures and dried by vacuum evaporation, then labeled for storage at -80 °C.

### 2.6 LC-MS/MS analysis

An EASY-nLC 1200 HPLC system (Thermo Fisher Scientific, USA) was used to separate the peptides. First, the peptides were acidified with 0.1% formic acid, their concentrations were determined by the BCA assay and then, they were diluted to 0.5 μg/μL with UA. Then, 1 μg of each peptides sample was loaded on the rap column (Acclaim PepMap^®^ 100, 75 μm×100 mm, 2 μm, nanoViper C18) at 0.3 μL/min (column flow rate) for 1 h (elution time). The elution gradient of mobile phase B was 5% to 40% (mobile phase A: 0.1% formic acid; mobile phase B: 89.9% acetonitrile). A Thermo Orbitrap Fusion Lumos Tribrid mass spectrometer (Thermo Fisher Scientific, USA) was used for analysis^[14]^. Survey MS scans were acquired by the Orbitrap in a 350-1550 m/z range with the resolution set to 120,000. For the MS/MS scan, the resolution was set at 30,000, and the HCD collision energy was chosen to be 30. Dynamic exclusion was employed with a 30-s window. Fifteen urine samples from three randomly selected experimental rats and three control rats at four time-points (days 3, 5, 7, and 11) were chosen for MS analysis. For each sample, two technical replicate analyses were performed.

### 2.7 Data analysis

All MS data were searched using Mascot Daemon software (version 2.5.1, Matrix Science, UK) with the SwissProt_2017_02 database (taxonomy: Rattus; containing 7992 sequences). The conditions included the following: trypsin digestion was selected, 2 sites of leaky cutting were allowed, cysteine was fixedly modified, methionine oxidation and protein N-terminal acetylation were mutagenic, peptide mass tolerance was set to 10 ppm and fragment mass tolerance was set to 0.05 Da. For statistical analyses that compared between the four-time points, one-way ANOVAs were performed. The differential proteins were screened with the following criteria: proteins with at least two unique peptides were allowed; fold change in increased group ≥1.5 and fold change in decreased group ≤0.67; and P <0.05 by independent sample t-test. Group differences resulting in P <0.05 were identified as statistically significant. All results are expressed as the mean ± standard deviation.

### 2.8 Functional annotation of the differential proteins

All differential proteins identified at the different time-points were analyzed by DAVID 6.8 (https://david.ncifcrf.gov/) and Ingenuity Pathway Analysis (IPA) to determine the functional annotation. The proteins were described in detail according to three aspects including biological process, cellular component and molecular function.

## 3. Results and Discussion

### 3.1 Body weight and histopathological characterization over time

There was a significant difference in body weight in the Walker 256 tumor-bearing rats over the seven days (Fig. 1). After W256 cell implantation, the average body weight of the tumor-bearing rats was lower than the controls, and reduced food and water intake was observed in the tumor-bearing rats. On day 16, a tumor-bearing rat died. All rats were sacrificed on day 18.

**Fig. 1.**
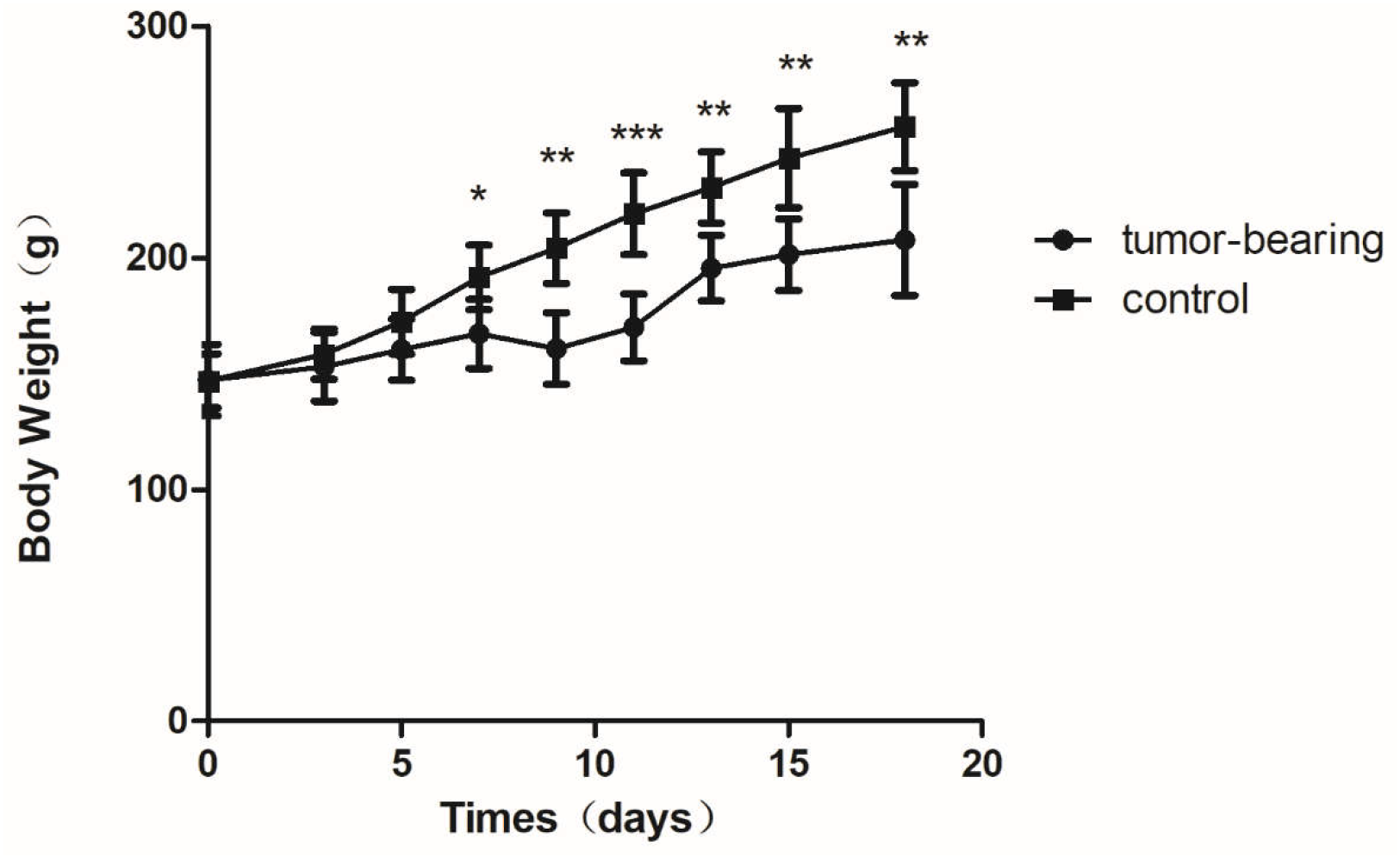
Body weights of Walker 256 tumor-bearing rats. The average body weight of the tumor group was significantly lower than that of the control group (n=6 rats in the tumor group and n=5 rats in the control group; * indicates p < 0.05; ** indicates p < 0.01; ***indicates p < 0.001).

H&E staining of HCC in the W256 rats showed that as the disease progressed to different stages, pathological changes increased. At day 3, the H&E staining showed that there were no obvious pathological changes. At day 7 and day 11, H&E staining observed carcinosarcoma cells under the microscope and the liver tissues revealed heterogeneously necrotizing tumors and liver tissue during tumor progression. At day 18, all the experimental rats exhibited fibrosis and the huge tumor that was viable while adjacent liver tissue was necrotizing (Fig. 2).

**Fig. 2.**
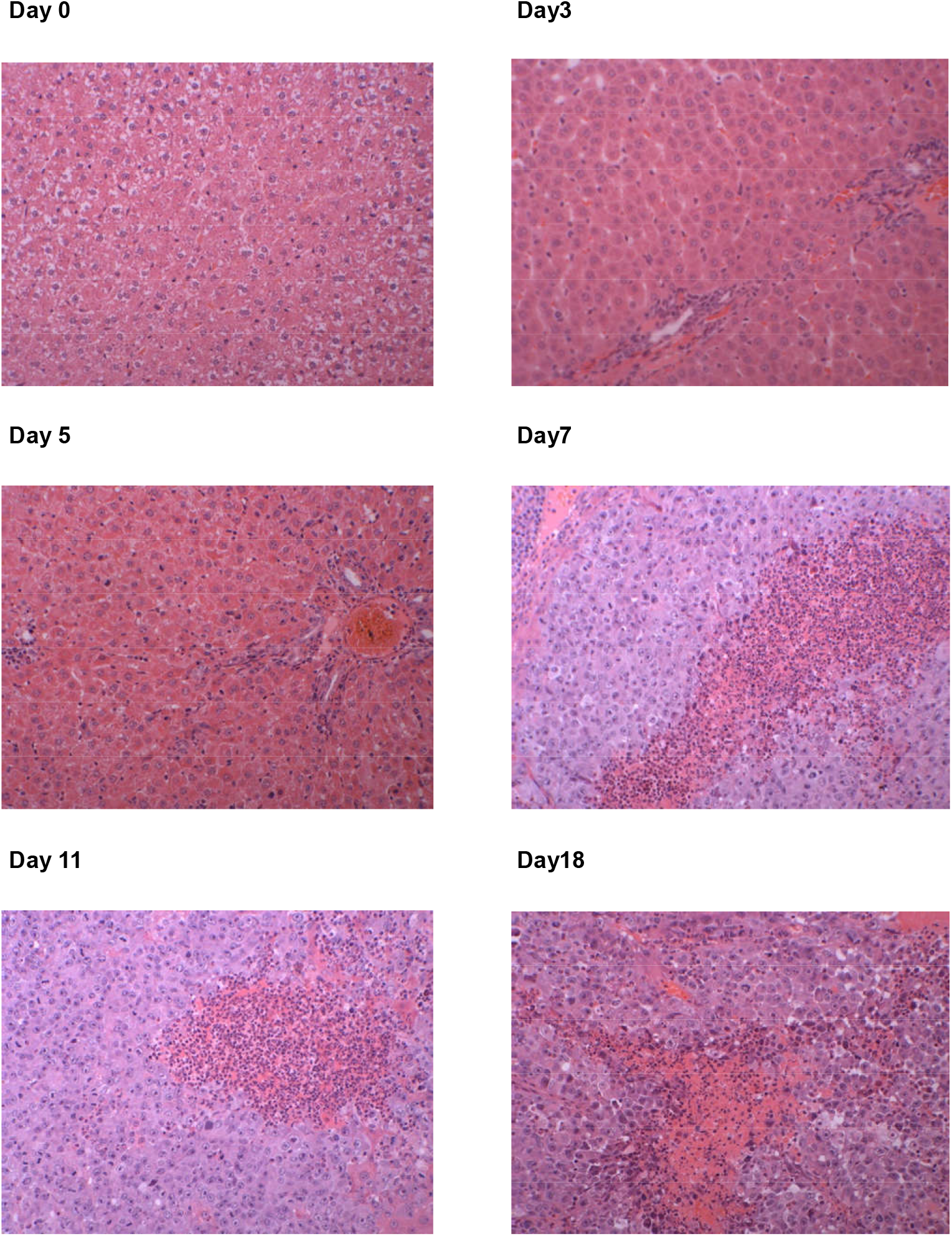
Histopathological characterization after implantation(200X).

### 3.2. Urine proteome changes in W256 model

To investigate how the urine proteome changes with tumor progression, urine samples from the four time-points (days 3, 5, 7, and 11) in three experimental rats and three control rats were chosen for MS analysis. In total, 663 urinary proteins were identified, and all proteins are listed in Table S1. Among these, there are 108 differential proteins and only 95 differential proteins that had human orthologs were identified that significantly changed in all rats (fold change ≥1.5 or ≤0.67, P < 0.05; Table 1).

**Table 1.**
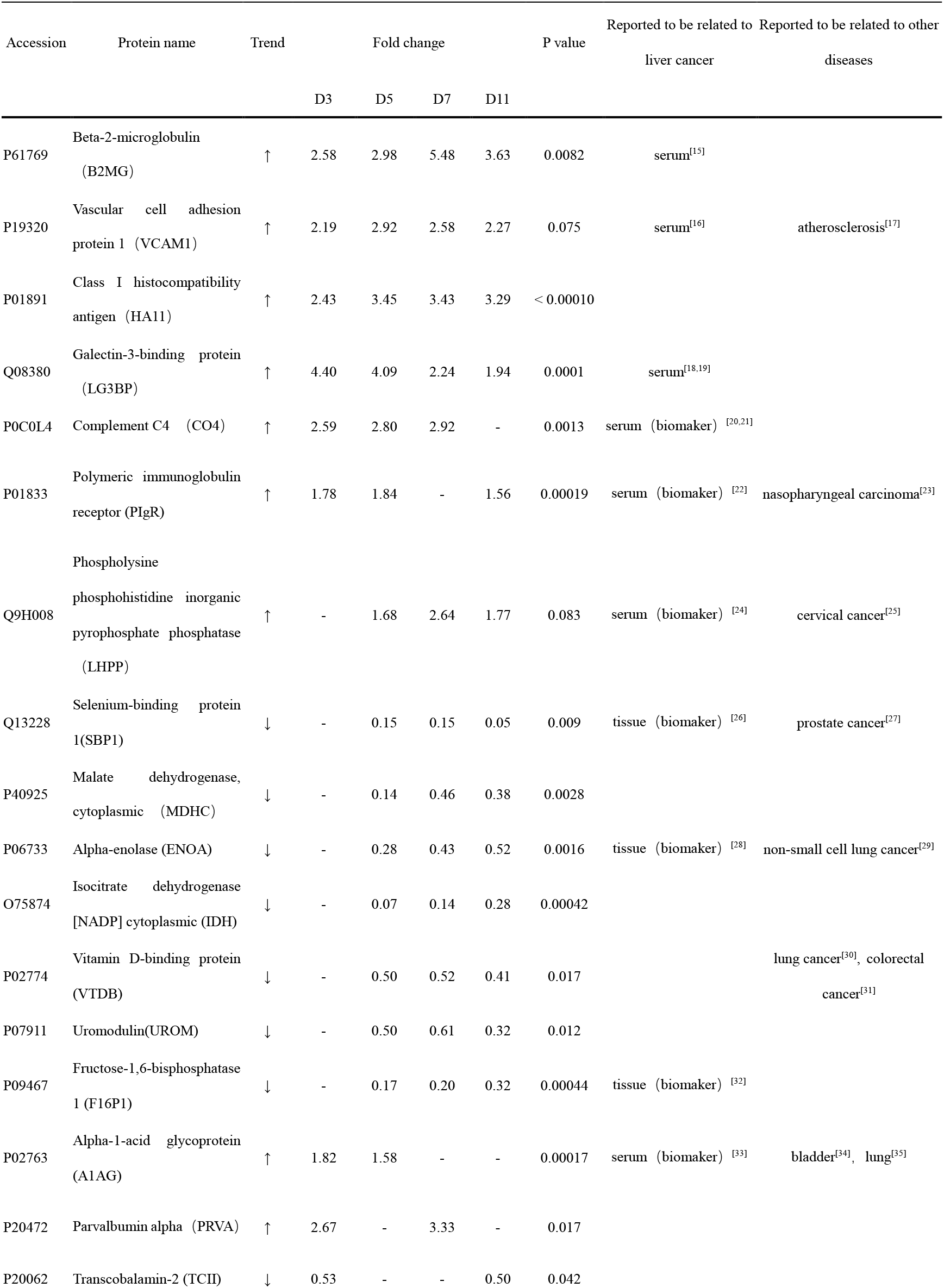
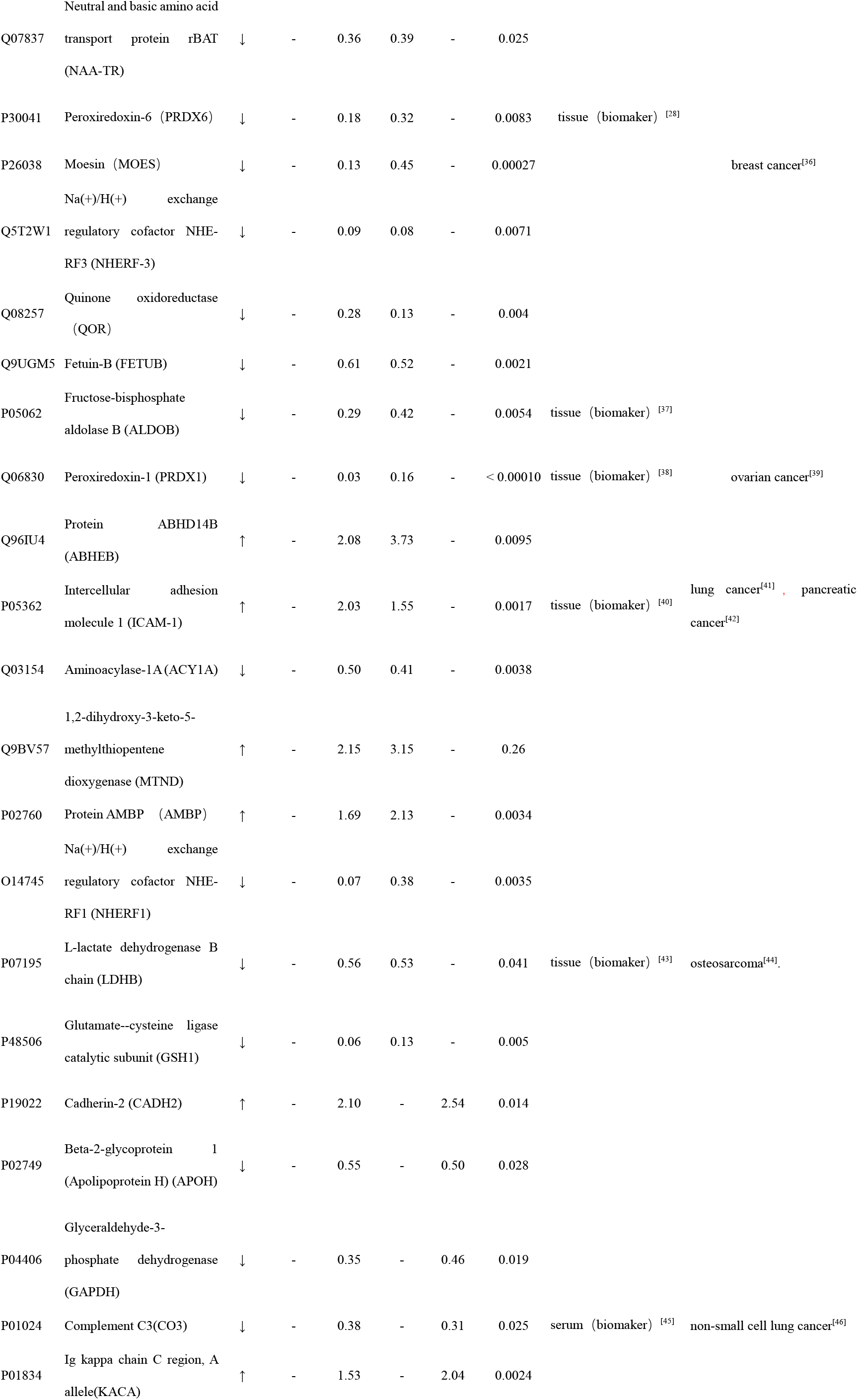
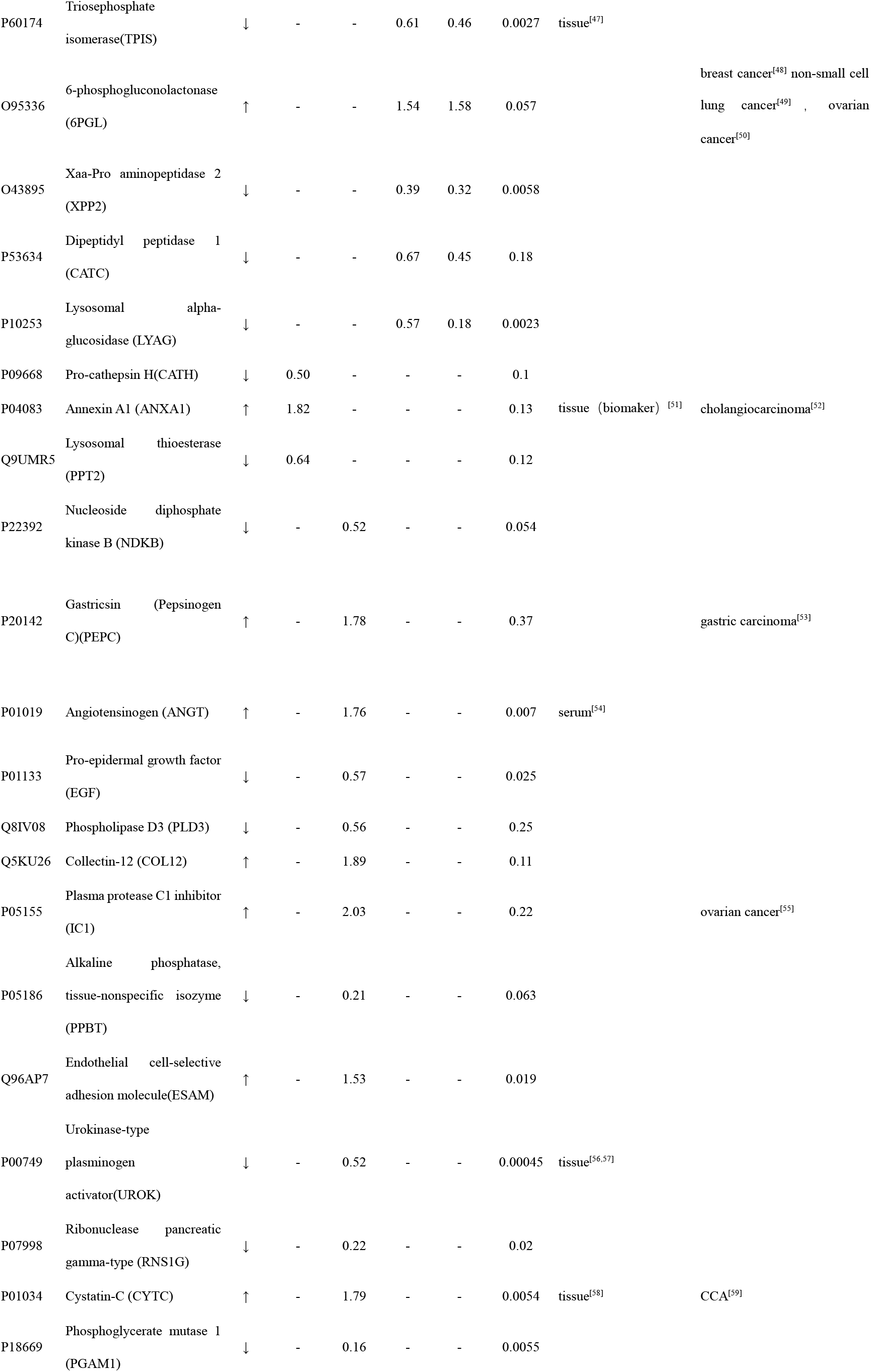
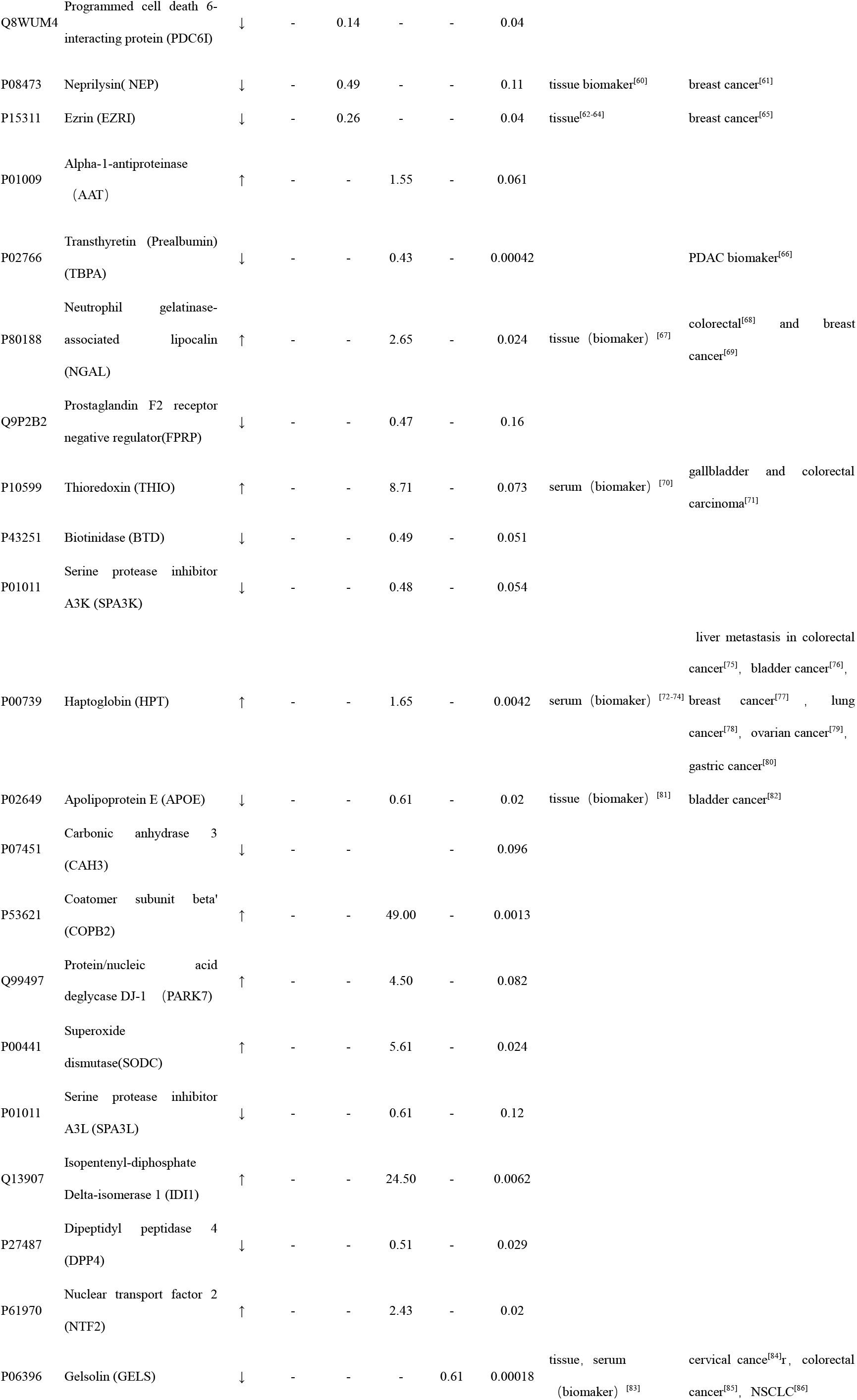
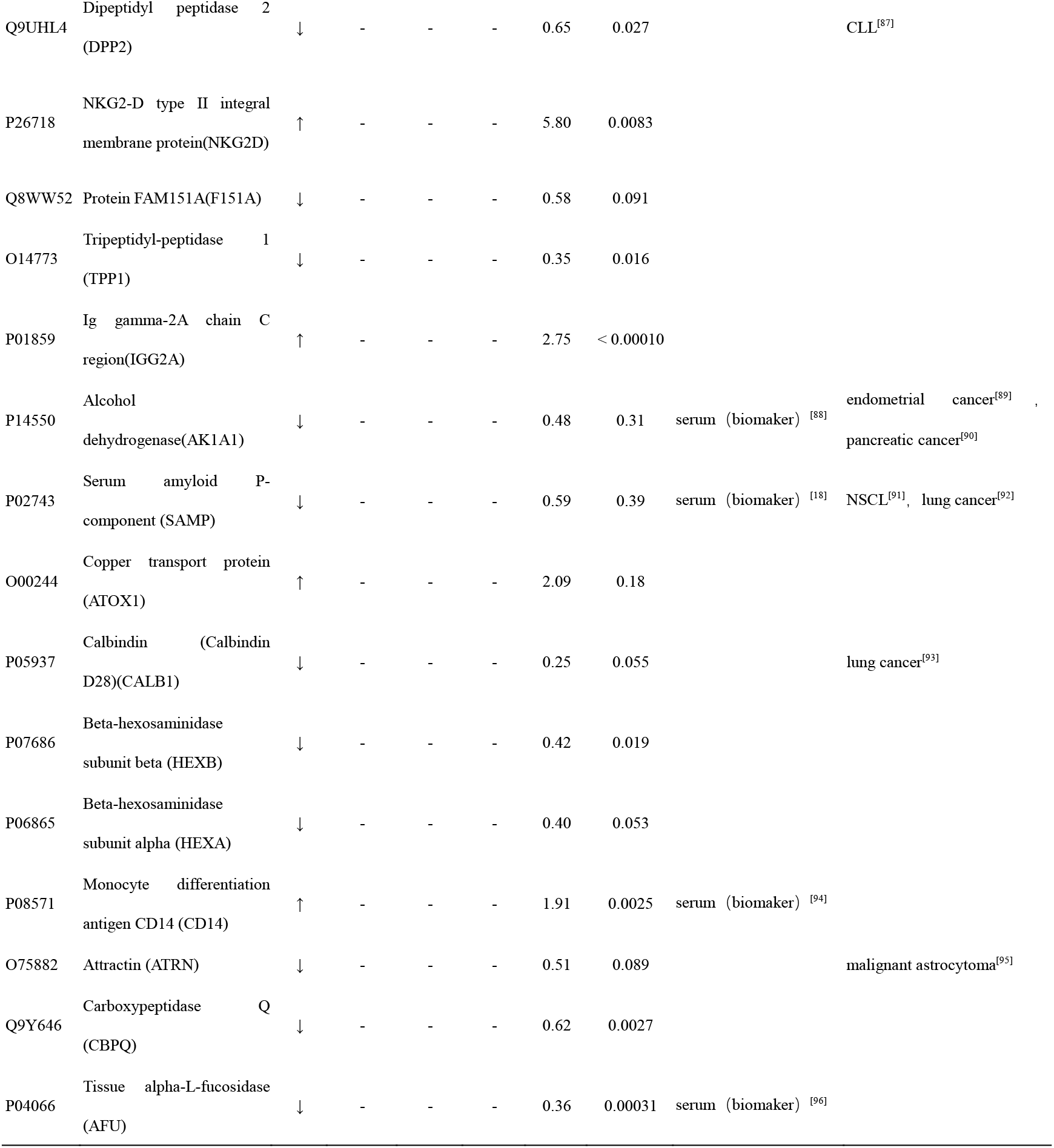
Differential urinary proteins in W256 model.

At day 3, twelve differential proteins, nine of which that increased and three of which that decreased, were identified. At day 5, fifty-two differential proteins, twenty of which that increased and thirty-two of which that decreased, were identified. At day 7, fifty-two differential proteins, twenty-two of which that increased and thirty of which that decreased, were identified. At day 11, forty differential proteins, thirteen of which that increased and twenty-seven of which that decreased, were identified. The details of the differential proteins are shown in Table S2. Four proteins (B2MG, VCAM1, HA11, LG3BP) were altered at all four time-points, and the trend was consistent at each time-point (Fig. 3B).

**Fig. 3.**
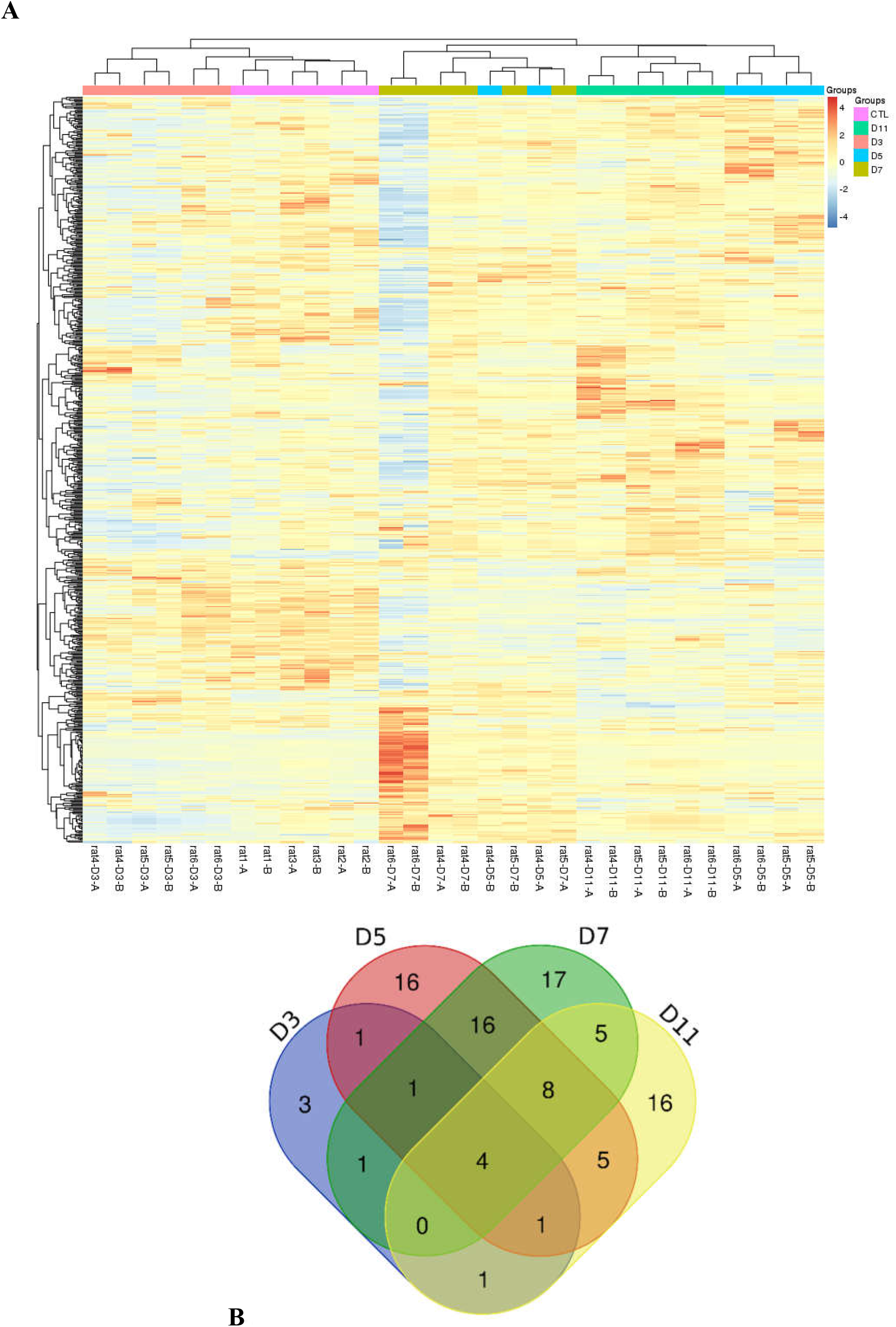
Statistical analysis of the urine proteome of W256 rats. A) Hierarchical clustering of the 663 proteins from the 15 samples (twelve subjects in the tumor-bearing group and three in the control group) at four time points. Lines represent proteins, and the colors correlate with their abundance (red indicates more abundant; blue indicates less abundant). B) The Venn diagram of the differential proteins identified at days 3, 5,7and 11.

### 3.3. Comparison of urinary proteins in different tumor models

The differential urinary proteins of four W256 tumor models (liver tumor model, lung metastasis model, intracerebral tumor model, and subcutaneous model) at all time points were compared and shown in Venn diagram (Fig.4).

**Fig. 4.**
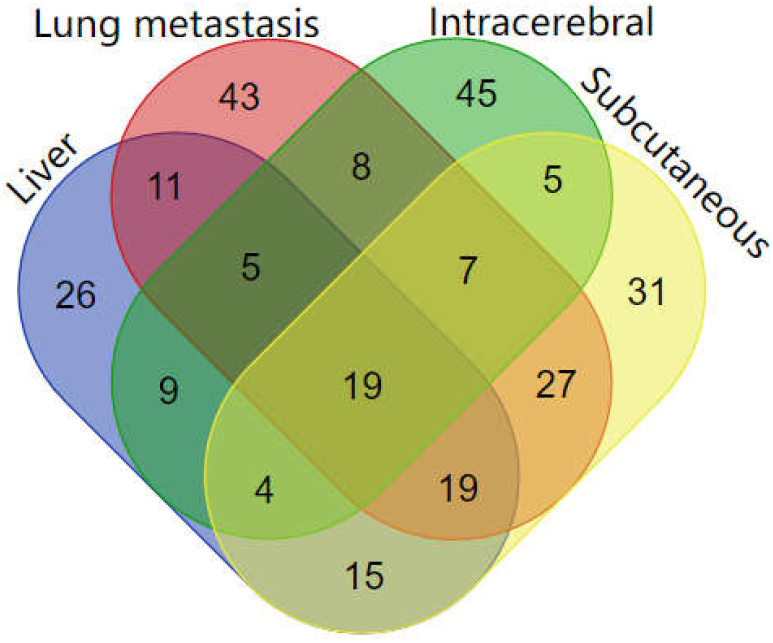
The overlapping differential proteins in urine samples of the four different W256 tumor models.

The result indicates that the same tumor cell grown in different organs can be reflected in differential urinary proteins. It can be seen from the Venn diagram that each model had a different number of unique differential urinary proteins. The 26, 43, 45, and 31 unique differential proteins were identified in the liver tumor model, the lung metastasis model, the intracerebral tumor model, and the subcutaneous model, respectively.

Twenty-one differential proteins had human orthologs were specially identified in W256 liver tumor model compared with the other three models. The comparison procedure is presented in Figure 5. Six of the 21 proteins (SAMP, LDHB, AFU, UROK, PRDX6, and PRDX1) had been reported to be associated with liver cancer. They were identified at the tumor development stages (D5, D7 and D13). (1) Serum amyloid P component (SAMP) is a candidate biomarker for HCC development in cirrhotic patients who were infected HCV^[18]^. (2) The expression of Lactate dehydrogenase B (LDHB) is a valuable prognostic biomarker for HCC. When the expression of LDHB becomes low, it suggests unfavorable survival outcomes^[43]^. (3) Alpha-l-fucosidase (AFU) is a lysosomal enzyme present in all mammalian cells and it has been proposed as a promising tumor marker since many studies reported increased AFU serum levels in patients with cirrhosis and HCC^[96]^. (4) Urokinase-type plasminogen activator(UROK) expression may be a potent therapeutic target of HCC^[97]^. (5) Peroxiredoxin 6 (PRDX6) may be a candidate biomarker for early HCC diagnosis^[28]^. (6) Peroxiredoxin 1 (PRDX1) is overexpressed in the tumor tissues of liver cancer and it also can predict poor prognosis for overall survival independently^[38]^.

**Fig. 5.**
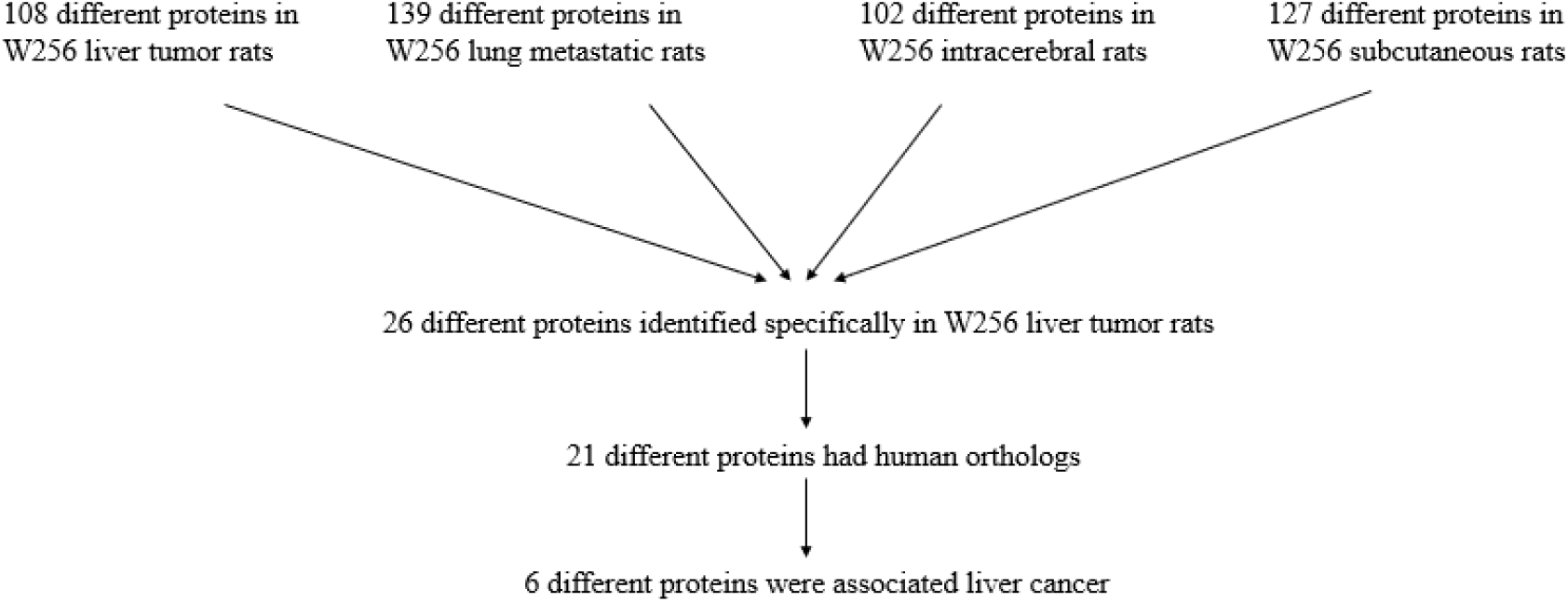
The comparison procedure of urinary proteins differentially expressed in the four models.

Among the 21 differential proteins, the remaining 15 proteins have not been reported as biomarkers of liver cancer. However, some of them were detectable at two time points. For example, it is reported that overexpression of cathepsin H(CATH) is related to several pathological states including carcinoma and melanoma^[98–100]^. Protein AMBP, a liver-specific precursor, is also a precursor of heme-binding protein that counteracts the disruption of free hemoglobin^[101,102]^. Cadherin-2 (CADH2) is called mesenchymal cadherin in carcinomas which play a role in epithelial-to-mesenchymal transition, and this process was considered to contribute to carcinoma progression^[103,104]^. Endothelial cell-selective adhesion molecule (ESAM), a member of the immunoglobulin receptor family, mediates homophilic interactions between endothelial cells. It suggests that ESAM has a special functional role in pathological angiogenic processes such as tumor growth^[105]^. Phosphoglycerate mutase 1 (PGAM1) is an important glycolytic enzyme that regulates many important biological processes, such as glycolysis, pentose phosphate pathway and serine biosynthesis in cancer cells^[106]^. The different functions of these proteins might be able to help distinguish the liver tumor and the tumors of other organs. These proteins may have potential for diagnosis and treatment of liver cancer in the future.

Among the overlapping proteins of these four models, it can be found that (1) 17 proteins had human orthologs can be detected in all models; (2) Most of the overlapping proteins reappear in more than two models in different combinations (Fig.4). One of the reasons for the appearance of these common proteins may be due to the same cells injected in all models; (3) Among the common proteins, 24 differential proteins had been reported to be associated with liver cancer, and some proteins had been identified as biomarkers in a variety of tumors; (4) The different combinations of the common proteins are also important to diagnosis. Because it is difficult to diagnose the type of tumor by using a single biomarker, thereby the panel of biomarkers is more accurate and reliable. Above all, the comparison results show that the growth of tumors in different organs has both commonality and individual difference. The urinary proteins have the potential to distinguish same tumor cell grown in different organs.

### 3.4 Functional analysis of differential proteins

In the W256 liver tumor model, the functional analysis of differential proteins at days 3, 5, 7 and 11 consisted of categorizing the biological processes, diseases, and functions using DAVID (Fig. 6). Ninety-eight differential proteins were annotated. In the biological processes, the innate immune response, retina homeostasis, response to drugs, negative regulation of endopeptidase activity, membrane to membrane docking, leukocyte cell-cell adhesion, complement activation classical pathway, and glycolytic process were significantly changed. At day 3, the innate immune response was the first to respond to the tumor cells. At day 5 and day 7, with the development of tumors in vivo, the glycolytic process, complement activation classical pathway, carbohydrate metabolic process, glutathione metabolic process, leukocyte adhesion, membrane to membrane docking, establishment of the endothelial barrier, and negative regulation of endopeptidase activity began to respond to the tumor changes. At day 11, the tumor grew further in the body, and the necrosis of some tissues caused phagocytosis recognition. The need for nutrients made some biological processes that were similar to the previous time-points continue. (Fig. 6A). In the cellular component, most of the differential proteins were in the extracellular exosome, extracellular space, MHC class I protein complex, blood microparticle, and extracellular region. They were changed in all the time-points. A small number of differential proteins come from organelles (Fig. 6B). In the molecular function category, endopeptidase inhibitor activity, identical protein binding, peroxiredoxin activity, and NAD binding were overrepresented. These biological processes were associated with neoplastic progression (Fig. 6C). From these analyses, it can be seen that these proteins caused these changes and their sources. This further confirmed that the changes in urinary protein come from the body’s response to the tumor cells. When the W256 cells entered the body, they caused an innate immune response. With the tumor development, the tumor cells further evaded or counterattacked the immune system through various pathways^[107]^.

**Fig. 6.**
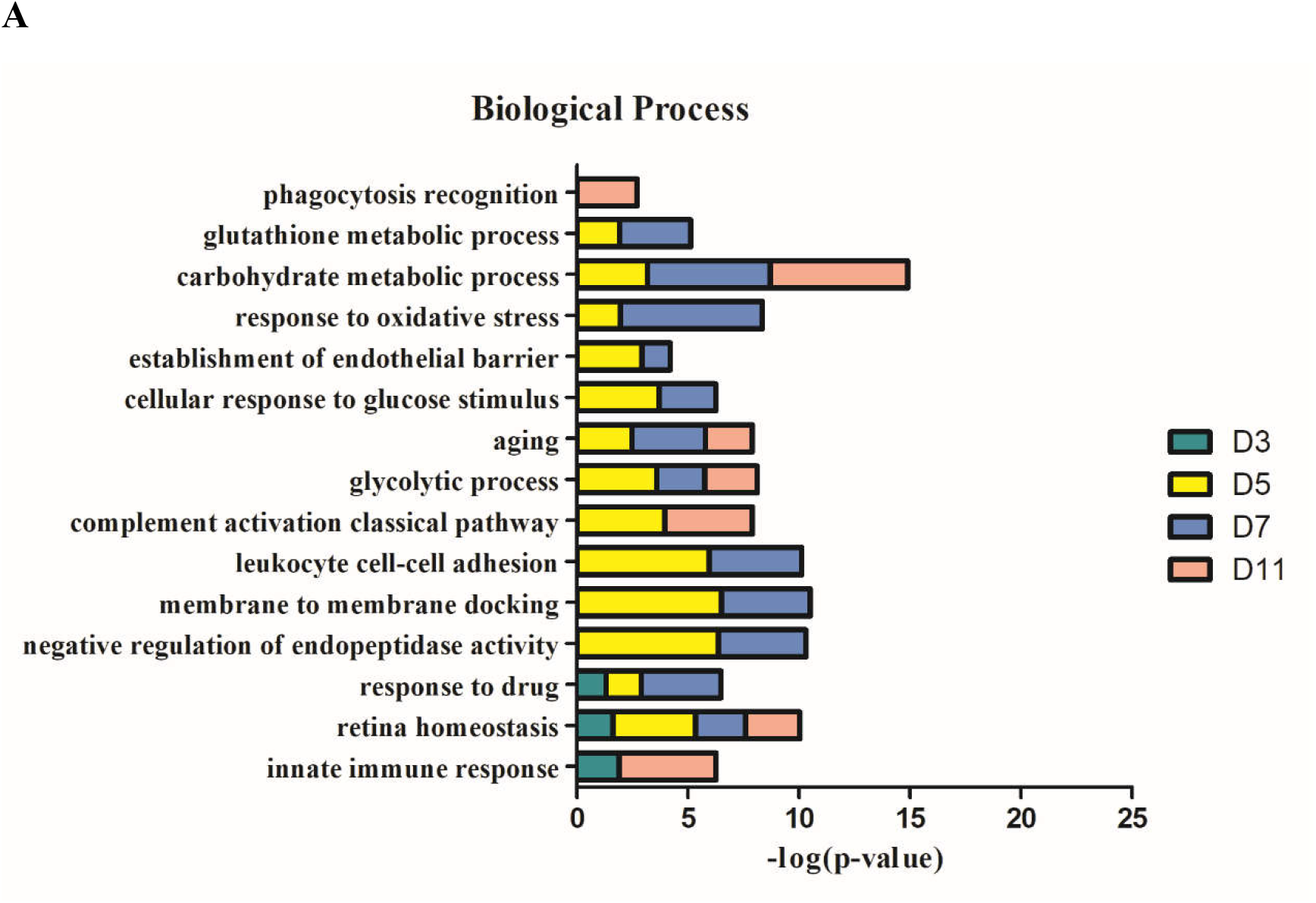
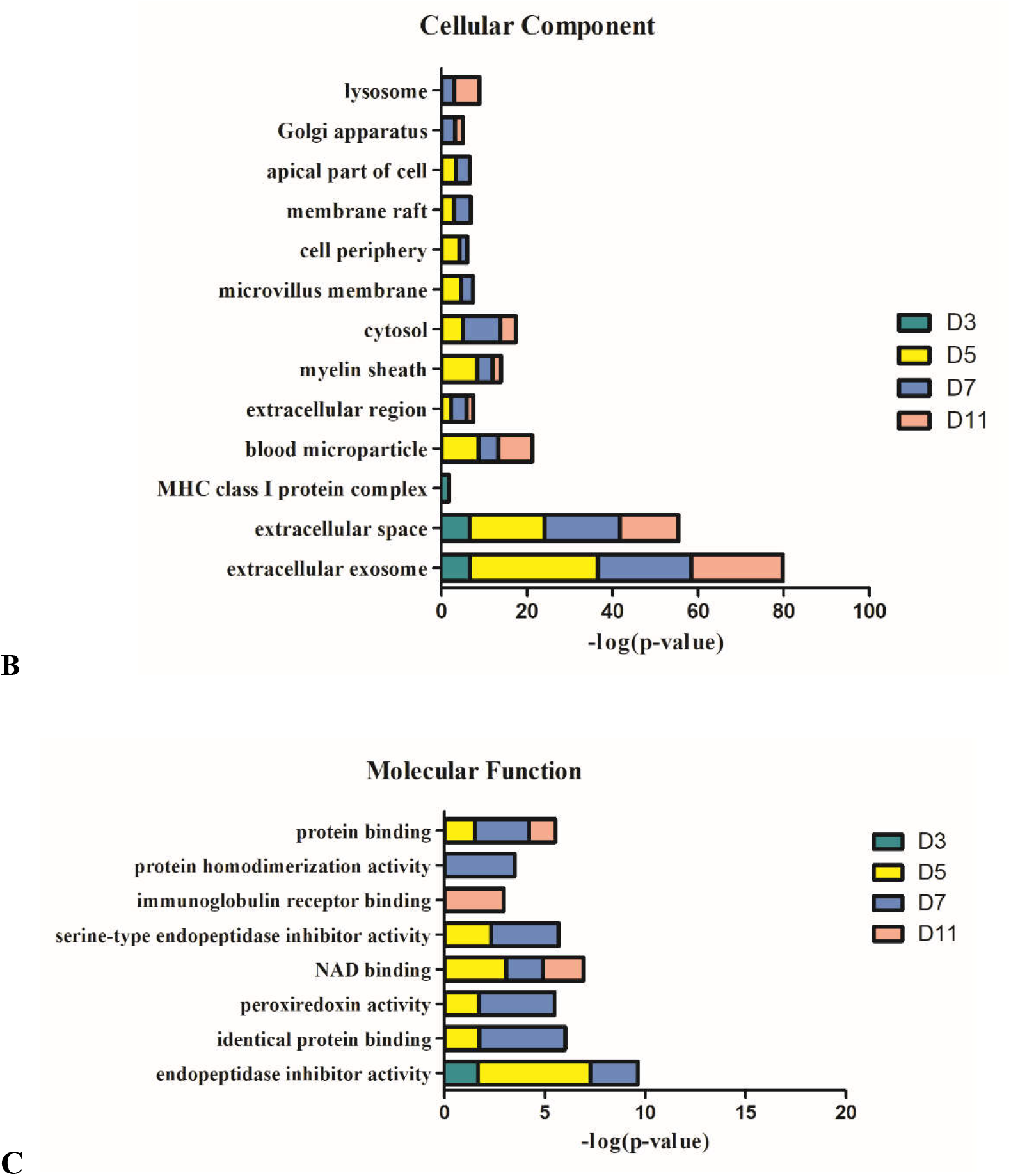
Functional analysis of differential proteins at days 3, 5, 7 and 11 in W256 model. A) Biological process; B) Cellular component; C) Molecular function.

The IPA analysis yielded several results. In the canonical pathway, FXR/RXR activation, gluconeogenesis I, glycolysis I, LXR/RXR activation, acute phase response signaling, allograft rejection signaling, phagosome maturation, OX40 signaling pathway, Cdc42 signaling and NRF2-mediated oxidative stress response showed the most marked changes. It was demonstrated that LXR/RXR activation, acute-phase response signaling, IL-12 signaling, production of nitric oxide and reactive oxygen species in macrophages, and the complement system were significantly enriched during tumor progression^[11]^. Gluconeogenesis I and glycolysis I also demonstrated the process of tumor development because of the increased glucose flux compared to normal tissue that is a common trait of human malignancies^[108]^. Some similar pathways have changed compared to previous experiments in which W256 cells were injected at other sites. The common pathways included acute-phase response signaling, LXR/RXR activation IL-12 signaling, and the complement system. The W256 tumor-bearing rat model has been previously used to study cancer-induced cachexia. Cachexia was characterized by weight loss after tumor cell inoculation^[109,110]^. This study also demonstrated results consistent with previous studies (Fig. 7).

**Fig. 7.**
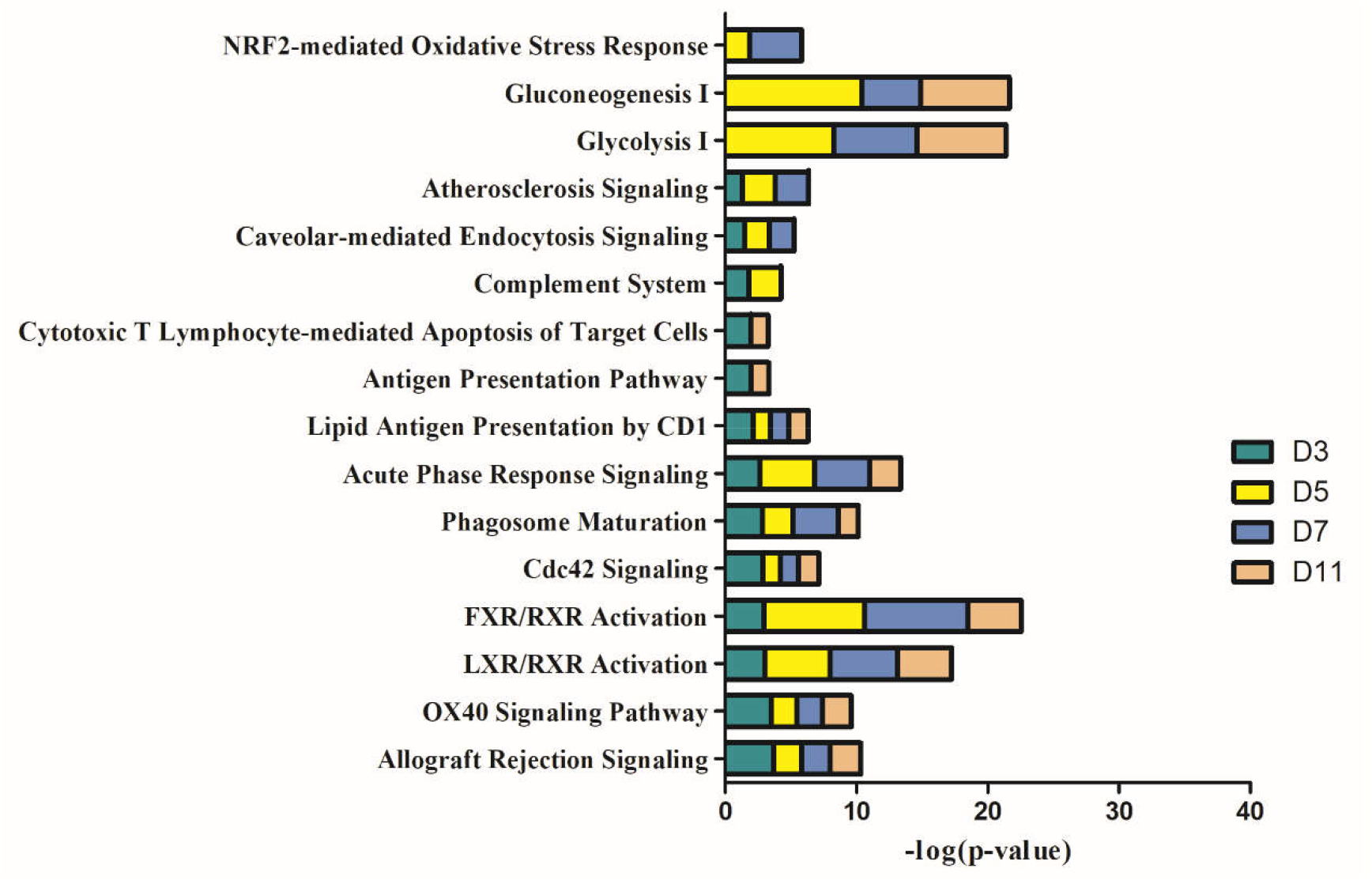
IPA analysis of differential proteins at days 3, 5, 7 and 11 in W256 model.

Figure 8 shows the comparison of biological processes of the liver tumor model with other models at early stages based on the data obtained from the studies of our laboratory published before^[11,111,112]^. The urinary proteins of different W256 models reflect different biological processes, suggesting that the biological processes of the same tumor cell grown in different organs may be different. In the early stages of all the models, the biological processes are very different. In the liver tumor model, the biological processes mainly reflect the immune response and metabolism. It may be due to that liver is a central organ for homeostasis and carries out a wide range of functions, including metabolism, glycogen storage, drug detoxification, production of various serum proteins, and bile secretion^[113]^. These biological processes are also associated with the functions of the liver. In the subcutaneous model, the biological processes are the primary response to various nutrients and ions. In the intracerebral tumor model, recognition and migration of cells in the biological processes are particularly significant. In the lung metastasis model, the biological processes include epithelial cell differentiation, regulation of immune system process, complement activation, classical pathway, ERK1 and ERK2 cascade, inflammatory response, etc. A large number of different early biological processes exist in the lung metastasis model for the reason that the differential proteins in the early stage are more than those of the other three models.

**Fig. 8.**
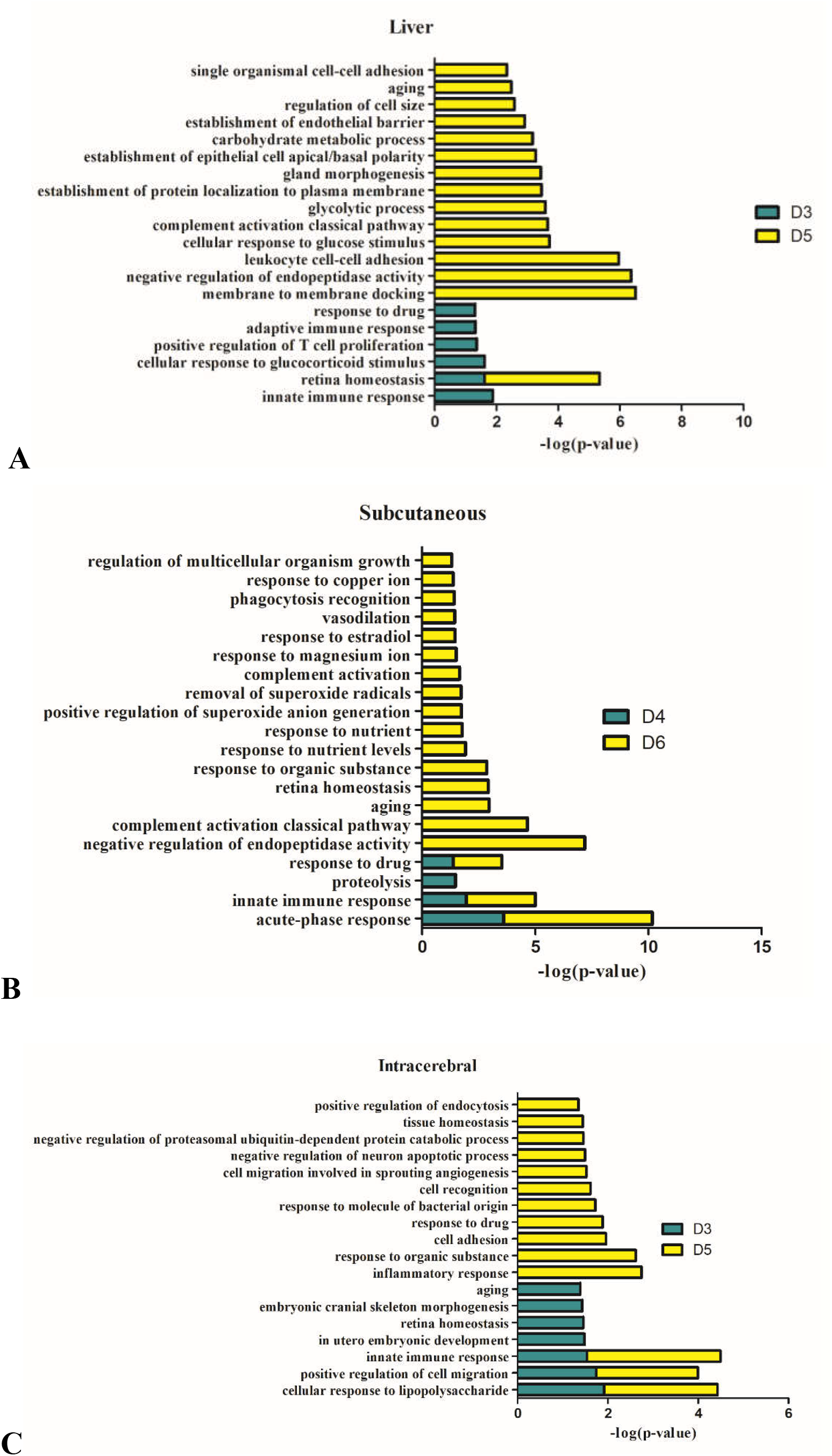
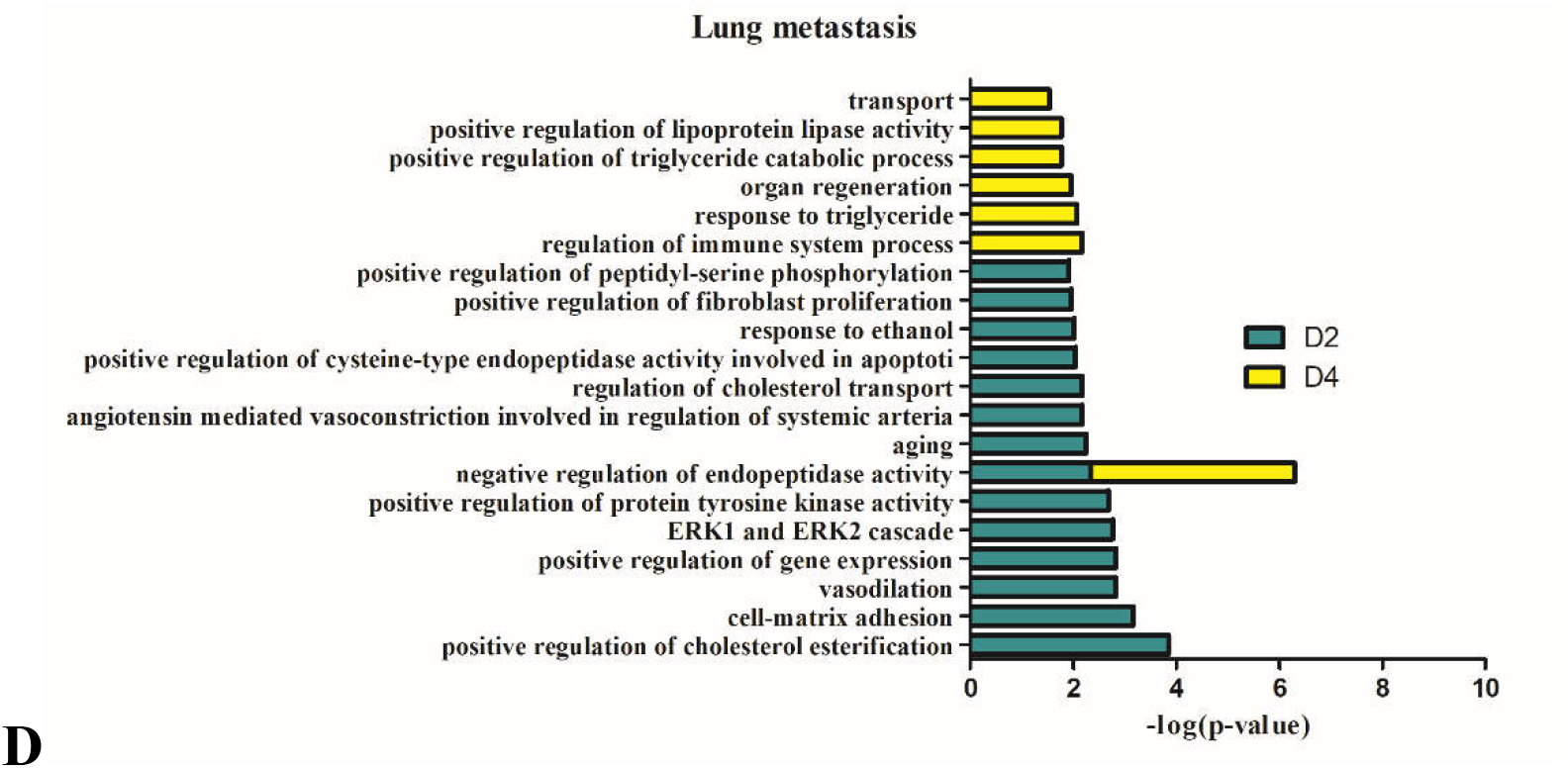
The analysis of the early stages of biological processes in different W256 models. A) The liver tumor model; B) the subcutaneous model; C) the intracerebral tumor model; D) the lung metastasis model. All the early biological processes are shown above. There are 34 early biological processes in the W256 lung metastasis model. For the convenience of comparison, the same number of biological processes as the W256 liver tumor model was selected according to p-value.

### 3.5. Analysis of differential proteins

In the W256 liver tumor model, the urinary proteins changed significantly after the tumor cells were implanted in the rats. Twelve proteins at day 3 changed significantly before the obvious pathological appearance, and fifty-two proteins changed at day 5. At day 3, nine of these proteins, LG3BP, PRVA, CO4, B2MG, HA11, VCAM1, ANXA1, A1AG, and PIGR, all were upregulated in liver cancer and other diseases in the serum or tissue. B2MG, VCAM1, HA11, and LG3BP were detected at all four time-points, and the other two proteins, CO4 (day 3, day 5 and day 7) and PIGR (day 3, day 5 and day 11), were found at three different time-points. A1AG continuously changed at day 3 and day 5. Galectin-3-binding protein (LG3BP) is a secreted glycoprotein that has an affinity for galectins and extracellular matrix proteins, and it can also interact and regulate cell adhesion^[15]^. It has been reported that LG3BP is expressed at high levels in various infectious and malignant diseases, such as HCV and HCC^[1,114]^. LG3BP was considered a poor prognosis biomarker in different types of malignancy^[115]^. Complement C4 (CO4) was reported as a potential biomarker to detect HCC in serum samples^[20,21]^. Beta-2-microglobulin (B2MG) may be used as a serum biomarker in HCV-related chronic liver diseases^[116]^. Vascular cell adhesion protein 1 (VCAM1) is closely related to the severity of the underlying liver disease^[16]^. Annexin A1 (ANXA1) is a member of the annexin superfamily proteins that can cause the pathological consequence and sequelae of many human diseases, and it can be used as a prognosis biomarker and a potential therapeutic target in HCC^[51]^. Alpha-1-acid glycoprotein (A1AG) was reported to be a potential biomarker in HCC patients^[33]^. Polymeric immunoglobulin receptor (PIGR) is one of the Fc receptor family members and is an important component of the mucosal immune system^[22]^. It has been proven to be a potential clinical target with the ability to promote cancer malignancy in HCC^[117]^. Of the other proteins, the fold change of parvalbumin alpha (PRVA) ranked second among the significantly upregulated proteins, and there were three downregulated proteins, lysosomal thioesterase (PPT2), transcobalamin-2 (TCII), pro-cathepsin H (CATH) and Class I histocompatibility antigen (HA11). Although there is a lack of reports associating them with liver cancer or other liver diseases, they may play an important role in the early stage of liver cancer.

At day 5, in addition to the proteins mentioned above, five of the upregulated proteins (ICAM1, CYTC, LHPP, A1AG, and ANGT) and ten of the downregulated proteins (LDHB, UROK, CO3, ALDOB, ENOA, EZRI, PRDX6, F16P1, SBP1, and PRDX1) were reported to be associated with liver cancer. Intercellular adhesion molecule 1 (ICAM1) belongs to the immunoglobulin superfamily and it has diagnostic significance in different kinds of HCC, such as AFP-negative or suspected HCC^[40]^. The expression of Cystatin C (CYTC) is higher in primary hepatic carcinoma than the control both at the tissue level and serum level^[58]^. There is a low rate of overall survival when the expression of histidine phosphatase (LHPP), a tumor suppressor, is reduced in patients with hepatocellular carcinoma^[24]^. Alpha-1-acid glycoprotein (A1AG) was found to be a novel biomarker to distinguish HCC plasma from control plasmas^[33]^. It has been reported that hepatoma is an angiotensinogen (ANGT)-producing tumor^[54]^. L-lactate dehydrogenase B chain (LDHB) is a valuable prognostic biomarker, and low expression suggests poor survival outcomes in HCC patients^[43]^. The Complement C3 (CO3) detection starts at a very early stage of tumor development, and it may represent a biomarker candidate for liver cancer^[45]^. The downregulation of fructose-bisphosphate aldolase B (ALDOB) suggests a poor prognosis, and it is a prognostic biomarker, especially at the early stage of HCC^[37]^. Alpha-enolase (ENOA) has been reported as biomarkers for HCC^[28]^. Ezrin (EZRI) expression can predict metastasis disease in human primary hepatocellular carcinoma tissue^[62]^. Fructose-1, 6-bisphosphatase 1 (F16P1) and selenium-binding protein 1 (SBP1) are considered potential biomarkers for prognosis in liver cancer^[26,32]^. Among the top-ranked proteins according to values of their fold change, for example, in the top ten of upregulation (MTND, CADH2, ABHEB, and IC1) and the top ten of downregulation (PGAM1, MDHC, PDC6I, MOES, NHRF3, IDHC, NHRF1, and GSH1), these unreported proteins also have great potential to be used to predict liver cancer. Although no relationship with liver cancer has been established, previous research suggests that they play important roles in other diseases (Table 1).

As the disease progressed, the number of proteins increased significantly at the last two time-points (day7 and day11). The pathological manifestations at these stages were also very obvious. Therefore, protein biomarker candidates were mainly selected at the time-points before pathological changes, especially those proteins that changed continuously, such as LG3BP, B2MG, CO4, VCAM1, HA11, A1AG, and PIGR. Among all differential proteins, several proteins were not only associated with liver cancers but also differentially changed in other cancers, suggesting that it is difficult to distinguish cancer types by relying on one or two protein markers. This may be related to the mechanism of tumor development.

Thus, this study suggests that a panel of urinary differential proteins is an ideal choice to improve the sensitivity and accuracy of early diagnosis for liver cancer.

## 4. Conclusion

Before the obvious pathological changes in the liver tumor model, the urinary differential proteins could be identified. Several differential proteins had been reported to be associated with liver cancer. These findings may provide important information for the early diagnosis of liver cancer. Additionally, the same tumor cell grown in different organs can be reflected in differential urinary proteins.

## Author contributions

Y.Z. and Y.H.Gao. conceived and designed the experiments. Y.Z. and Y.F.Gao. performed the experiments. Y.Z. analyzed the data and wrote the manuscript. All authors approved the final manuscript.

## Acknowledgements

This research was supported by the National Key Research and Development Program of China (2016 YFC 1306300), Key Basic Research Program of the Ministry of Science and Technology of China (2013FY114100), Beijing Natural Science Foundation (7173264, 7172076), Beijing cooperative construction project (110651103), Beijing Normal University (11100704), and Peking Union Medical College Hospital (2016-2.27).

